# Canonical NOTCH signaling controls the early progenitor state and emergence of the medullary epithelial lineage in fetal thymus development

**DOI:** 10.1101/600833

**Authors:** Dong Liu, Anastasia I. Kousa, Kathy E. O’Neill, Francois Guillemot, Martyna Popis, Alison M. Farley, Simon R. Tomlinson, Svetlana Ulyanchenko, Philip A. Seymour, Palle Serup, Ute Koch, Freddy Radtke, C. Clare Blackburn

## Abstract

Thymus function depends on the epithelial compartment of the thymic stroma. Cortical thymic epithelial cells (cTECs) regulate T cell lineage commitment and positive selection, while medullary (m) TECs impose central tolerance on the T cell repertoire. During thymus organogenesis, these functionally distinct sub-lineages are thought to arise from a common thymic epithelial progenitor cell (TEPC). The mechanisms controlling cTEC and mTEC production from the common TEPC are not however understood. Here, we show that emergence of the earliest mTEC lineage-restricted progenitors requires active NOTCH signaling in progenitor TEC and that, once specified, further mTEC development is NOTCH-independent. In addition, we demonstrate that persistent NOTCH activity favors maintenance of undifferentiated TEPC at the expense of cTEC differentiation. Finally, we uncover a direct interaction between NOTCH and FOXN1, the master regulator of TEC differentiation. These data establish NOTCH as a potent regulator of TEPC and mTEC fate during fetal thymus development and are thus of high relevance to strategies aimed at generating/regenerating functional thymic tissue *in vitro* and *in vivo*.

## Introduction

In the thymus, thymic epithelial cells (TECs) are the essential stromal component required for T lymphocyte development (Manley et al., 2011; Ritter and Boyd, 1993). Two functionally distinct TEC subsets, cortical (c) TECs and medullary (m) TECs, exist and are found in the cortex and the medulla of the organ respectively. Thymocytes migrate in a highly stereotypical fashion to encounter cTECs and mTECs sequentially as T cell differentiation and repertoire selection proceeds (Anderson and Takahama, 2012; Klein et al., 2014).

cTECs and mTECs originate from endodermal progenitor cells (thymic epithelial progenitor cells; TEPCs), that are present in the thymic primordium during its initial generation from the third pharyngeal pouches (3PPs) (Gordon et al., 2004; Le Douarin and Jotereau, 1975; Rossi et al., 2006). Several studies have shown that, during development, both cTECs and mTECs arise from cells expressing markers associated with mature cTECs, including CD205 and β5t (Baik et al., 2013; Ohigashi et al., 2013), while clonal analyses have shown that a bipotent TEPC can exist *in vivo* (Bleul et al., 2006; Rossi et al., 2006). Based on these observations, a serial progression model of TEC differentiation has been proposed (Alves et al., 2014). This suggests that fetal TEPCs, which exist as a transient population, exhibit features associated with the cTEC lineage and that additional cues are required for mTEC specification from this common TEPC. Identification of cTEC-restricted sub-lineage specific progenitor TECs in the fetal thymus has proved elusive, due to the shared expression of surface antigens between this presumptive cell type and the presumptive common TEPC (Alves et al., 2014; Baik et al., 2013; Shakib et al., 2009), although cTEC-restricted progenitors clearly exist in the postnatal thymus (Ulyanchenko et al., 2016). In contrast, the presence of mTEC-restricted progenitors has been detected from day 13.5 of embryonic development (E13.5) (Rodewald et al., 2001). In the fetal thymus, these mTEC progenitors are characterised by expression of Claudin3/4 and SSEA1 (Hamazaki et al., 2007; Sekai et al., 2014). Receptors leading to the activation of NFκB pathway, including LTβR and RANK, are known to regulate the proliferation and maturation of mTEC through crosstalk with T cells and tissue inducer cells (Boehm et al., 2003; Hikosaka et al., 2008; Rossi et al., 2007) and, recently, a hierarchy of intermediate progenitors specific for the mTEC sub-lineage has been proposed based on genetic analysis of NFκB pathway components (Akiyama et al., 2016; Baik et al., 2016). Additionally, HDAC3 has emerged as an essential regulator of mTEC differentiation (Goldfarb et al., 2016), and a role for STAT3 signaling has been demonstrated in mTEC expansion and maintenance (Lomada et al., 2016; Satoh et al., 2016). Despite these advances, the molecular mechanisms governing the emergence of the earliest cTEC- and mTEC-restricted cells in thymic organogenesis are not yet understood (Hamazaki et al., 2007).

NOTCH-signaling has been extensively studied in the context of thymocyte development (Shah and Zuniga-Pflucker, 2014), and is also implicated as a regulator of TECs. Mice lacking the Notch ligand JAGGED 2 showed reduced medullary areas (Jiang et al., 1998a; Jiang et al., 1998b), while B cells overexpressing another Notch ligand, Delta like 1 (DLL1), induced organized medullary areas in a reaggregate fetal thymic organ culture (RFTOC) system (Masuda et al., 2009). In contrast, in adult thymic epithelium NOTCH activity appeared to reside in cTECs, while its TEC-specific overexpression reduced TEC cellularity and led to an imbalance between mature and immature mTECs, suggesting that NOTCH signaling might inhibit mTEC lineage development (Goldfarb et al., 2016). Overall, these results suggest that NOTCH has complex effects in TECs, but the stage(s) at and mechanism(s) through which NOTCH influences TEC development have not yet been determined.

We have addressed the role of NOTCH signaling in early TEC differentiation using loss- and gain-of-function analyses. Our data establish, via genetic ablation of NOTCH signaling in TECs using *Foxn1^Cre^;Rbpj^fl/fl^* and *Foxa2^Cre^;dnMAML* mice, and via fetal thymic organ culture (FTOC) in the presence of NOTCH-inhibitor, that NOTCH signaling is required for specification of the mTEC lineage. They further demonstrate that the initial sensitivity of mTEC to NOTCH is restricted to a time-window prior to E16.5, and that NOTCH is required earlier than RANK-mediated signaling in mTEC development. Finally, they show that NOTCH signaling is permissive rather than instructive for mTEC specification, since TEC-specific overexpression of Notch Intracellular Domain (NICD) in fetal TEC dictated an undifferentiated TEPC phenotype rather than uniform adoption of mTEC characteristics. Collectively, our data establish NOTCH as a potent regulator of TEPC and mTEC fate during fetal thymus development.

## Results

### Early fetal mTECs exhibit high NOTCH activity

To begin to understand how NOTCH signaling affects thymus development, we first investigated the expression of NOTCH ligands and receptors in TEC during early organogenesis, via RT-qPCR of E10.5 3PP cells and defined E12.5 to E14.5 TEC populations separated on the basis of EPCAM, PLET1 and UEA1 expression as appropriate (Fig.1; for gating strategies see Supplementary Fig. 1). *Notch1, Notch2, Notch3, Jagged 1* (*Jag1*) and *Delta like 4* (*Dll4*), but no other NOTCH receptors and ligands, were expressed throughout this time period (Fig. 1). *Notch1* and *Notch2* were significantly enriched in E14.5 UEA1^+^ mTECs compared to all other populations examined. *Notch3* and *Jag1* were more highly expressed in PLET1^+^ and UEA1^+^ TEC than in other TEC subpopulations, with *Notch3* being most highly expressed at E10.5 (Fig. 1A). Of the NOTCH target genes examined, *Hes1* and *Heyl* showed similar expression patterns to *Notch3* from E12.5. In contrast and as anticipated, the highly expressed NOTCH ligand and direct FOXN1 target *Dll4* was initiated at E12.5. At E13.5 and E14.5, *Dll4* was more highly expressed in PLET1^−^ than in PLET1^+^ TEC and in cTECs than mTECs, respectively, consistent with the *Foxn1* expression pattern (Fig. 1A). At the protein level, at E13.5 NOTCH1 was enriched in UEA1^+^ TECs (NOTCH1^+^ among UEA1^+^, 51.5% ± 8.4%) compared to UEA1^−^ TECs (24.7% ± 10.4%) (Fig. 1B). NOTCH2 and JAG1 were also coexpressed with UEA1 at E14.5, while NOTCH3 was more broadly expressed (Fig. 1C). Furthermore, analysis of the CBF:H2B-Venus mouse line, which reports NOTCH signaling (Nowotschin et al., 2013), indicated ongoing or recent NOTCH activity in half of E14.5 UEA1^+^CD205^−^ mTECs compared to only a small minority of cells in the CD205^+^UEA1^−^ ‘cTEC’ population (Fig. 1D). Collectively, these data show that the earliest TECs experience high levels of NOTCH signaling, while early mTECs remain competent to receive further NOTCH signals.

**Figure 1.**
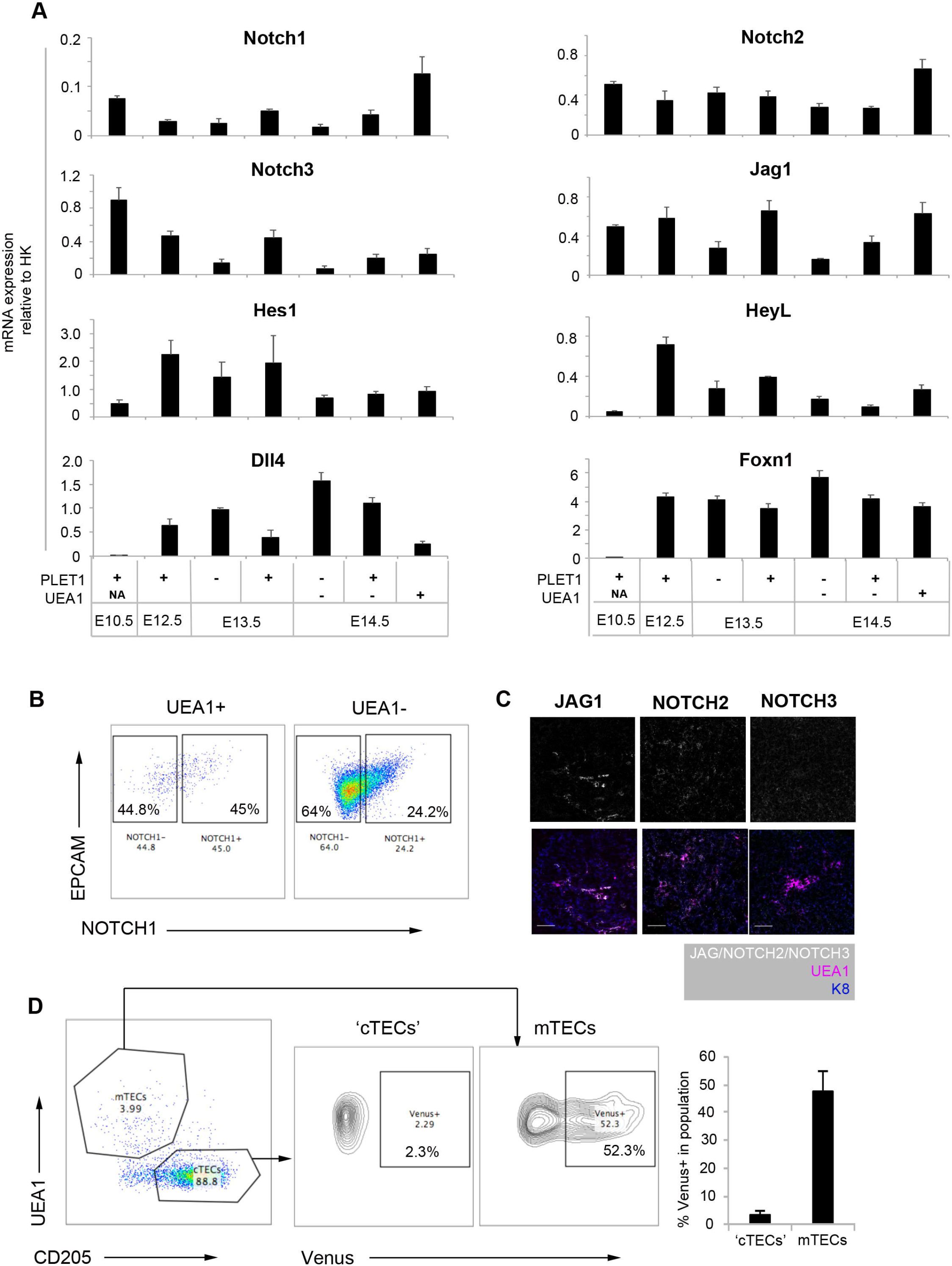
Expression of NOTCH pathway components in thymus organogenesis. Dynamic mRNA expression of NOTCH receptors, ligands and targets between E10.5 and E14.5. (B) Typical flow cytometry plots of NOTCH1 expression in E13.5 TECs, split by expression of UEA1. (C) Single images of JAG1, NOTCH2 and NOTCH3, and co-staining with mTEC marker UEA1 and epithelial marker K8 on sections of E14.5 thymus primordium. Scale Bar=50μm. (D) Left: Representative profile of E14.5 CBF1:H2B-Venus thymi, gated on EPCAM^+^ epithelial cells. Cell suspension was stained with the mTEC marker UEA1 and the cTEC/progenitor (‘cTEC’) marker CD205. Middle: Proportion of ‘cTECs’ and mTECs showing the expression of Venus. Right: Quantitation of the percentage of Venus expression in E14.5 ‘cTEC’ and mTEC populations. ***Data collection:*** (A) n = 3 (all genes at E10.5 and E14.5, Notch 3, Jag1, Heyl, Dll4 at E12.5 and E13.5) or 6 (Notch 1, Notch 2, Hes1 and Foxn1 at E12.5 and E13.5). In each case, n represents RNA obtained from pooled cells of the phenotype stated from an independent litter of embryos. (B) Plots shown are representative of n=3. each ‘n’ represents cells obtained from pooled thymi from an individual wild type litter. (C) n = 3 independent immunohistochemistry analyses. (D) n=4. Each ‘n’ is an independent E14.5 embryo from the same CBF1:Venus x C57BL6 litter; genotypes were retrospectively confirmed. ***Statistics:*** All error bars show mean±SD.

### NOTCH signaling is required for mTEC development

We next addressed the role of NOTCH in TEC development, by crossing *Foxn1^Cre^* mice (Gordon et al., 2007) to the *Rbpj^fl/fl^* conditional knock out mouse line (Han et al., 2002). This generated mice in which RBP-Jκ was absent from all TEC and at least some cutaneous epithelial cells, rendering these cells unable to respond to NOTCH signaling (Han et al., 2002). The recombination efficiency of *Foxn1^Cre^* was close to 100% in E14.5 EPCAM^+^ TECs when tested using a silent GFP (sGFP) reporter (Gilchrist et al., 2003) (Supplementary Fig. 2), and genotyping indicated complete deletion of *Rbpj* in total TECs purified from 4-week-old *Foxn1^Cre^; RBPJ^fl/fl^* thymi (Supplementary Fig. 2). Having validated the *Foxn1^Cre^; RBPJ^fl/fl^* model (called *Rbpj* cKO herein), we next analyzed the effect of loss of RBPJ on the postnatal thymus. This revealed a significant proportional and numerical decrease in mTECs in both male and female *Rbpj* cKO mice at two weeks of age (Fig. 2A), with cTEC numbers unaffected (Fig. 2B). The decrease in mTEC numbers reflected reduced numbers of MHC Class II^hi^ (mTEC^hi^) and MHC Class II^lo^ (mTEC^lo^) TEC in males, and of mTEC^hi^ in females (Fig. 2B). This phenotype normalized by eight weeks of age, after which a second loss of mTEC was observed (Fig. 2C-E). No other RBP-Jκ-dependent thymic phenotypes were observed: T cell development in the *Rbpj* cKO mice was not blocked at any stage, and no difference in any of the intrathymic Treg precursor or Treg populations (CD25^−^FOXP3^+^, CD25^+^FOXP3^−^, CD25^+^FOXP3^+^)(Lio and Hsieh, 2008; Tai et al., 2013) was detected versus controls (Fig. 2F, Supplementary Fig. 2C). Thus, the thymic phenotype in the *Rbpj* cKO model appeared TEC-specific and affected mTEC but not cTEC.

**Figure 2.**
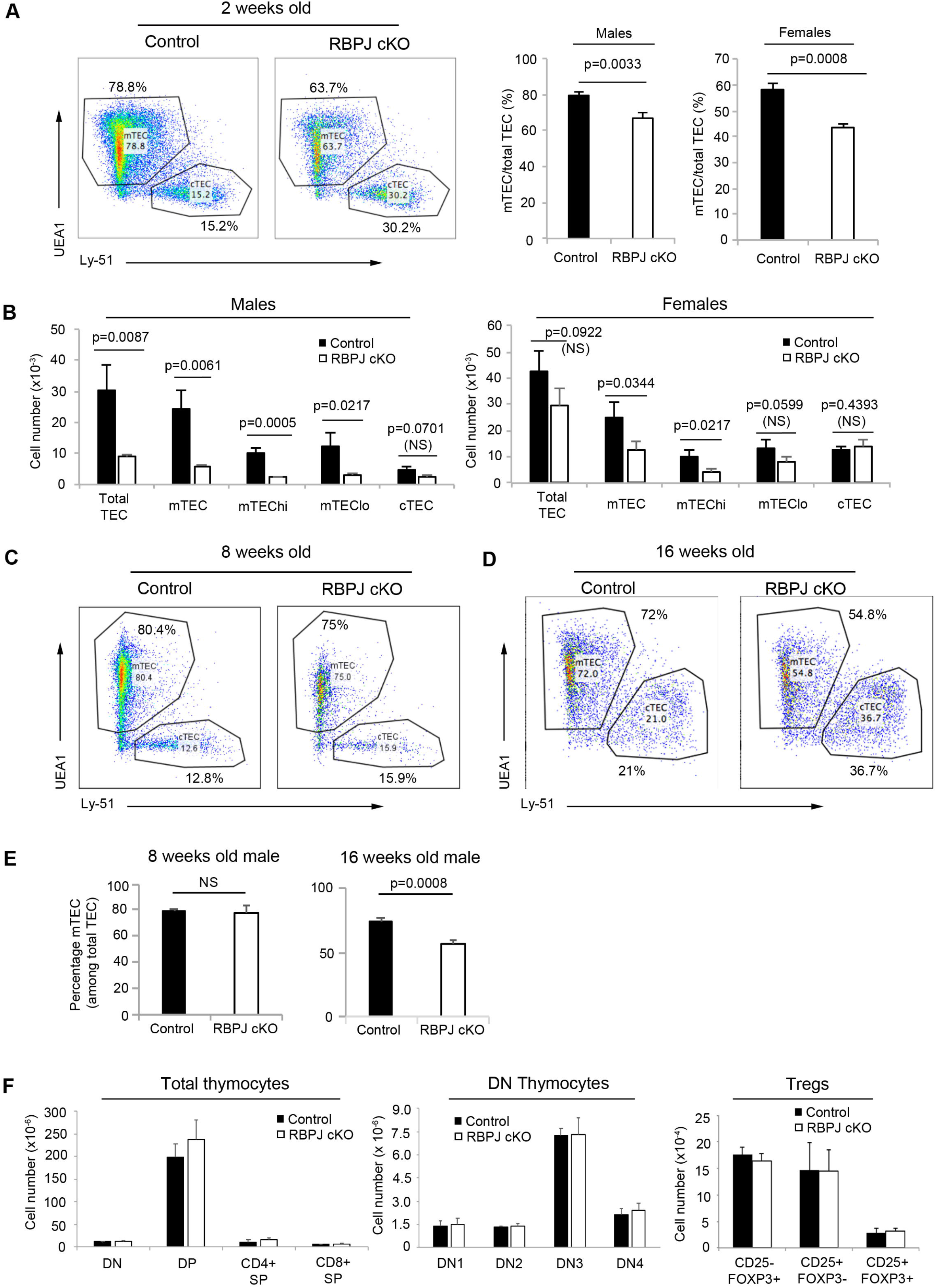
Loss of *Rbpj* leads to a proportional and numerical reduction of mTECs in postnatal thymus. (A) Left: Proportions of mTECs (UEA1^+^) and cTECs (Ly51^+^) in 2 weeks-old males. Right: mTEC proportions in 2 weeks-old males and females. (B) Absolute cell count of total TEC and sub-populations in 2 weeks-old males (left) and females (right). (C, D, E) Proportion of mTECs and cTECs in 8 (D) and 16 (E) week-old males. 8 weeks mTEC - WT 79.97±1.28, *Rbpj* cKO 78.27±4.07. (F) Left and middle: Absolute numbers of thymocyte subsets in 2 weeks-old females. Right: Absolute numbers of CD25^−^FOXP3^−^, CD25^+^FOXP3^−^, and CD25+FOXP3+ Tregs in 2 weeks-old males. Tregs were pre-gated as CD4^+^TCRβ^hi^CCR6^−^. ***Data collection:*** (A, B, F) n=3 cKO and 3 littermate control mice for male and female. (C-E) 8 weeks, n=3 cKO and 3 littermate control male mice, 16 weeks n=3 cKO and 3 littermate control male mice from 3 independent litters; results were confirmed in females (not shown). ***Statistics:*** All error bars show mean±SD. (F) p values in pairwise comparisons were calculated with two-tailed t-test.

### Temporal requirement for NOTCH signaling in mTEC development

To determine whether the *Rbpj* cKO mTEC phenotype arose postnatally or during development, we then analyzed E14.5 control and *Rbpj* cKO thymi using markers characteristic of developing mTEC and cTEC. Fewer K14^+^ and UEA1^+^ presumptive mTEC were present in E14.5 cKO thymi than in littermate controls (Fig. 3A). This indicated that the medullary phenotype was evident by E14.5, three days after the onset of Cre expression*/Rbpj* deletion, establishing that NOTCH signaling is required during emergence of mTEC lineage cells and pointing to a potential role in TEC progenitors.

**Figure 3.**
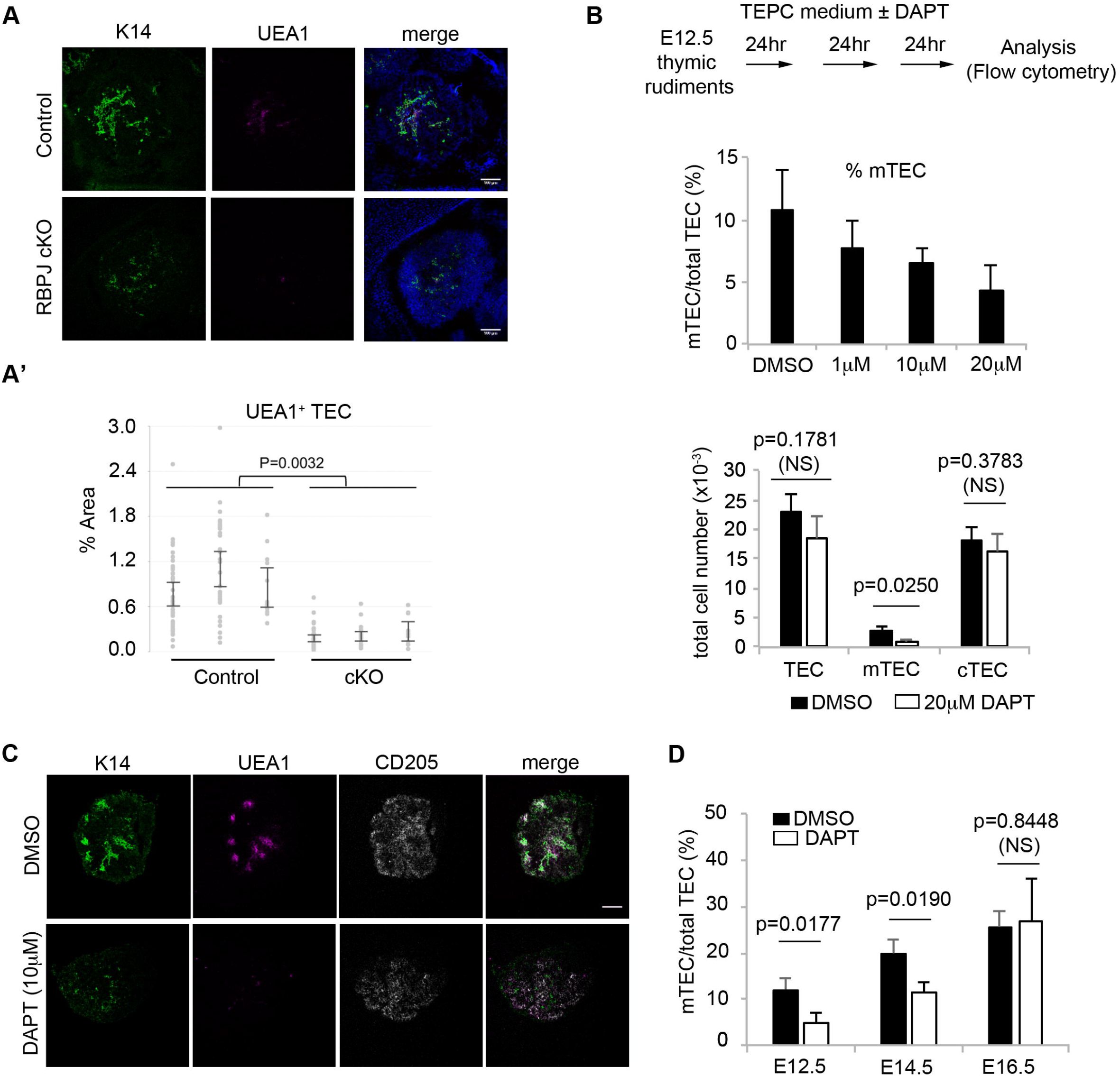
Loss of NOTCH signaling responsiveness by E14.5 results in diminished mTEC production. **(A)** The deficiency of mTECs in *Rbpj* cKO can be detected at E14.5. Representative transverse sections of the embryo showing the thymus primordium stained with the mTEC markers K14 and UEA1 and with DAPI to reveal nuclei. (**A**’) Proportion of pixels in the thymic section (within the outline of DAPI) stained positive for UEA1. (B) Proportion (top) and numbers (bottom) of mTEC in E12.5 explant cultures at different concentrations of DAPT. No significance difference was observed for total TECs or cTECs. (C) Representative sections of E12.5 thymic rudiments after FTOC in control condition or 10μM DAPT, stained with mTEC markers K14 and UEA1, and cTEC marker CD205. Error bar=100μm. (D) Proportion of mTECs after 3 days of FTOC in control condition or DAPT. The age of thymi at the onset of culture is indicated. Only data using the highest concentration of DAPT (20μM for E12.5 and E14.5; 50μM for E16.5) were collated. ***Data Collection:*** (A, A’) UEA1 images are representative of data collected from 3 cKO and 3 littermate control embryos from 3 separate litters. K14 images are representative of data collected from 4 cKO and 4 control embryos from 4 separate litters. Embryos were snap frozen in OCT. cKO and control embryos were selected for analysis following genotyping. (A’) Each data point represents a section, and each mean value represents the reconstruction of all thymus-containing sections of an embryo. (B-D) Each ‘n’ (i.e. each data point in the graphs) represents the proportion or number of cells from each of 9 fetal thymic lobes cultured under the given conditions. Each experiment was repeated three times. Several litters of wildtype embryos were used for each experiment. ***Statistics:*** (A’) Error bars show 95% confidence interval. (F) Error bars show mean±SD. p values in pairwise comparisons were calculated with two-tailed t-test.

This phenotype was independently confirmed using γ-secretase inhibition (DAPT) in fetal thymic organ culture (FTOC) (Fig. 3B). Addition of DAPT to the culture medium in FTOC cultured at the air-liquid interface (Ueno et al., 2005) (E15.5 or older) or submerged (E14.5 and younger) (Supplementary Figs. 3-5) resulted in down-regulation of NOTCH receptors, ligands and targets (Supplementary Fig. 3B). Moreover, E12.5 primordia cultured for three days in the presence of DAPT contained significantly fewer UEA1^+^ mTEC than controls (Fig. 3B). DAPT treatment had no effect on cTEC numbers or overall cellularity in this model (Fig. 3B). Control explants contained medulla-like foci that co-expressed K14 and UEA1 (Fig. 3C), similar to E15.5 thymic primordia (Fig. 3A) (Rodewald et al., 2001), while a substantial reduction in K14 and UEA1 expressing cells was observed in the DAPT-treated explants (Fig. 3C). These effects could not be attributed to treatment-induced apoptosis or decreased proliferation, since the proportions of Caspase^+^ and Ki67^+^ mTEC were not significantly affected at the concentration of DAPT used (20μM) (Supplementary Fig 4A, B). To examine the time-dependence of NOTCH signaling in early mTEC development, we extended these analyses by culturing E14.5 and E16.5 fetal thymi ± DAPT for three days. mTEC numbers in cultured E14.5 thymic primordia were significantly reduced in the presence of DAPT (Fig. 3D). However, in contrast, the percentage of mTECs in E16.5 thymi after three days of culture was unaffected (Fig. 3D). Collectively, NOTCH signaling regulates mTEC development during early thymus organogenesis in a restricted time window up to and including E15.5 but prior to E16.5.

### NOTCH acts prior to NF-κB signaling to regulate mTEC lineage progression

The NF-κB pathway ligands (Receptor activator of nuclear factor kappa-B ligand [RANKL], lymphotoxin beta and CD40L) are potent regulators of mTEC development and thymic lympho-epithelial crosstalk (Boehm et al., 2003; Hikosaka et al., 2008). Of these only RANKL stimulates both proliferation of mTEC and upregulation of the autoimmune regulator (*Aire*). Recent studies have shown that the expression of RANK receptor and hence responsiveness to RANKL stimulation increases with increasing maturation of mTEC progenitors (Akiyama et al., 2016; Baik et al., 2016; Mouri et al., 2011). To map the requirement for NOTCH-relative to RANK-signaling, we cultured E15.5 FTOC in the presence of deoxyguanosine (dGuo) to deplete T cells, and then in the presence of RANKL and the presence or absence of DAPT. RANKL elicited a proportional increase in mTEC as expected, (Fig. 4A) and co-treatment with DAPT mildly attenuated this response (Fig 4A). This suggested that NOTCH and NF-κB might independently regulate different aspects of mTEC development or could act sequentially in the same developmental pathway. To discriminate between these possibilities, we cultured E15.5 *Rbpj* cKO and littermate control thymi in dGuo-FTOC conditions with or without RANKL. Consistent with the data shown in Figures 2 and 3, some mTEC progenitors arose in the *Foxn1^Cre^Rbpj^fl/fl^* model. Culture of *Rbpj* cKO thymi in RANKL resulted in an approximately threefold proportional increase in mTEC versus unstimulated cKOs and these mTECs displayed a more mature phenotype (MHCII^+^) than controls, indicating that once generated, these mTEC progenitors respond normally to RANK. Nevertheless, in RANKL-stimulated *Rbpj* cKO thymi the proportion of mTEC was substantially lower than that in RANKL-stimulated wild-type controls (Fig 4B), placing the requirement for NOTCH signaling developmentally upstream of that for RANK. These data, together with those in Figure 3D, demonstrate a limited window for NOTCH regulation of mTEC progenitor emergence and establish that NOTCH signaling acts at an earlier stage than NF-κB signaling to regulate the number of mTEC progenitors. They further indicate that once mTEC progenitors are specified, NOTCH is dispensable for mTEC differentiation.

**Figure 4.**
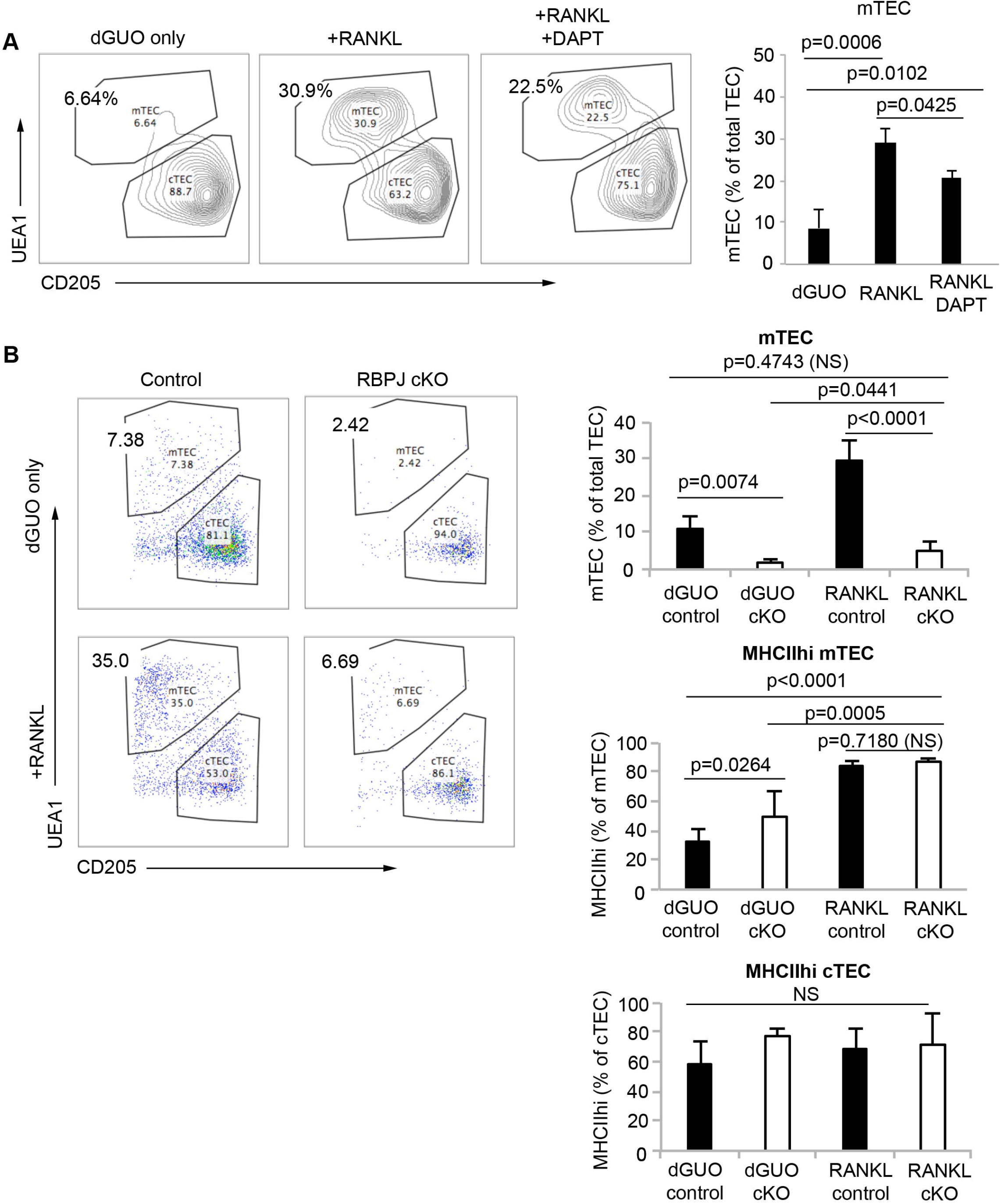
NOTCH is required prior to NFκB signaling in mTEC development. (A) (Left) Representative plots showing the proportion of UEA1^+^ mTECs and CD205^+^ cTECs after 3 days in FTOC. E15.5 thymi were used and the concentration of DAPT was 50μM. (Right) Quantitation of mTEC proportions after FTOC. The clear effect of DAPT observed on *in vitro* development of E15.5 lobes cultured at the air-liquid surface also indicates its effective penetration into thymic lobes in these culture conditions. (B) (Top) Representative plots showing the proportion of UEA1^+^ mTECs and CD205^+^ cTECs after 3 days in FTOC. E15.5 thymi were used. The condition and genotype are labelled on vertical and horizontal axes respectively. (Bottom) Quantitation of the percentage of mTECs, and the percentage of MHCII^+^ cells in mTEC and cTEC populations. ***Data Collection:*** (A) n=3, where each n represents an independent experiment in which 3 wildtype E15.5 thymic lobes were treated in each condition. (B) E15.5 thymi from three litters from a *Foxn1^Cre^;Rbpj^FL/+^ x Rbpj^FL/FL^* cross were cultured either with or without RANKL. Litters were obtained and cultured on different days. Genotypes for each embryo were determined retrospectively. No samples were excluded from the analysis. For each condition, each n represents the thymic lobes from a single embryo; dGuo control, n=6; dGuo cKO, n=5; RANKL control, n=5; RANKL cKO, n=4. ***Statistics:*** All error bars show mean±SD. p values were calculated with one-way ANOVA test (two tailed).

### NOTCH signaling is required for specification of the mTEC lineage

The above data would be consistent with NOTCH-regulation of mTEC specification, mTEC progenitor expansion, or both. The *Foxn1^Cre^;Rbpj* cKO model results in deletion of *Rbpj* from around E12.0, with subsequent loss of RBP-Jκ function depending on protein turnover and cell division time. The emergence of mTEC progenitors has however been suggested by phenotypic studies to occur independently of FOXN1, possibly at least as early as E10.5 (Hamazaki et al., 2007; Nowell et al., 2011), and therefore the presence of reduced numbers rather than total loss of mTEC progenitors in this model may reflect the relatively late timing of RBP-Jκ deletion. Furthermore, *Rbpj* mRNA is expressed at only very low levels in E12.5 TEC (not shown), suggesting that NOTCH-mediated effects should occur prior to this time-point. To discriminate between the above possibilities, we therefore determined the effect of blocking NOTCH signaling in TEC at or prior to mTEC and cTEC lineage divergence. For this we generated mice in which NOTCH-mediated transcription is blocked in the developing endoderm before E9.5, by crossing the *Foxa2^T2AiCre^* line with mice carrying the inducible dominant negative Mastermind allele *Rosa26^loxp-STOP-loxp-dnMAML-IRES-eGFP^* allele (Horn et al., 2012; Maillard et al., 2004). This *Foxa2^T2AiCre^;Rosa26^loxp-STOP-loxp-dnMAML-IRES-eGFP^* model (referred to herein as dnMAML) provides a stronger and earlier block of NOTCH activity than that in *Foxn1^Cre^;Rbpj^fl/fl^* (i.e. *Rbpj* cKO) mice.

E14.5 dnMAML thymi appeared smaller than controls but contained thymocytes and endothelial networks (Supplementary Fig 7; controls were aged-matched *Foxa2^T2iCre^;Gt(ROSA)26So^rtm1(EYFP)Cos^* embryos). At E14.5, CLDN3^+^ TECs are mTEC-lineage restricted and contain cells with long-term mTEC reconstituting activity (Hamazaki et al., 2007; Sekai et al., 2014). Crucially, at E14.5 this CLDN3^+^ TEC population was completely/almost completely absent from dnMAML thymi (mean 8.6-fold reduction in dnMAML thymi, with some thymi exhibiting a complete loss) (Fig. 5A,B,D; note that the CLDN3 staining seen in Fig. 5B is restricted to endothelial cells). Similarly, the number of K14^+^ mTEC was reduced dramatically in E14.5 dnMAML thymi versus littermate controls (Fig. 5C; note that the reduction is more pronounced than that in E14.5 *Rbpj* cKO thymi). A profound effect on mTEC development was also evident in E16.5 dnMAML thymi, with some thymi containing no K14^+^, UEA1^+^ or AIRE^+^ mTEC and others containing one or two foci staining with one or more of these markers (Fig. 5E-G; 3.7-fold decrease in K14^+^ area; 6.6-fold numerical reduction in AIRE^+^ mTECs). These data indicate that blockade of NOTCH-mediated transcription prior to E9.5 results in a near complete block in mTEC progenitor production, effectively resulting in a ‘medulla-less’ thymus.

**Figure 5.**
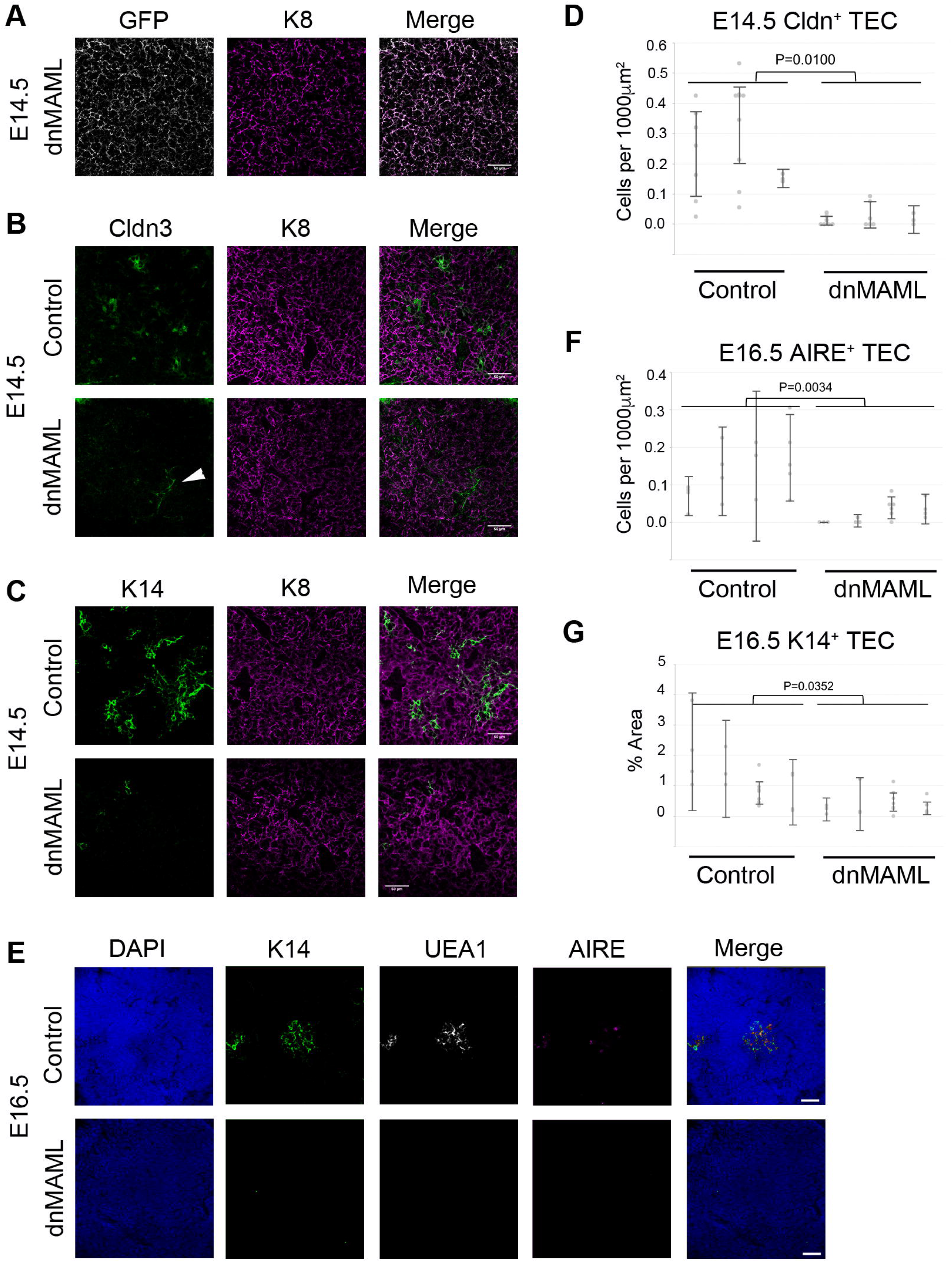
NOTCH signaling is an essential mediator of mTEC specification. (A) Representative E14.5 dnMAML section, showing the overlap between GFP (recombined cells) and K8 (TECs). Scale bar=50μm. (B, C) Representative images of E14.5 control and dnMAML sections stained for mTEC progenitor marker Claudin3 (CLDN73; B), mTEC marker K14 (C) and epithelial marker K8. Scale bar=50μm. (D) Quantification of the number of CLDN3^+^ TECs in E14.5 control and dnMAML thymi. Note that some weakly stained CLDN3^+^ cells co-localized with endothelial marker CD31 (white arrowhead in B, and see Supplementary Figure 7A), hence for quantification only CLDN3^+^K8^+^ double positive cells were counted. (E) Representative images of E16.5 sections stained for DAPI, UEA1, K14 and AIRE. Scale bar=50μm. (F-G) Quantification of the number of AIRE^+^ mTECs (F) and area of K14^+^ staining (area of marker [over positive threshold]/area of thymus defined by DAPI staining) (G) in E16.5 control and dnMAML thymi. ***Data Collection:*** *Foxa2^T2iCre^;Rosa26^loxp-STOP-loxp-dnMAML-IRES-eGFP^* and *Foxa2^T2iCre^;Gt(ROSA)26Sor^tm1(EYFP)Cos^* (control) embryos were collected at E14.5 and E16.5. Samples analyzed were littermates. (D,F,G) Each data point represents a section. Mean values from several images from the same embryo were used for statistics. E14.5, n=3; E16.5, n=4. ***Statistics:*** Error bars show 95% confidence interval. p values were calculated with the twotailed unpaired t-test.

This conclusion was supported by explant culture of E10.5 3PP. Initial validation of the culture system (Supplementary Fig. 5) showed that during five days of culture, E10.5 3PP explants undergo morphogenesis, differentiation and self-organization consistent with continuing development of the thymus primordium (Supplementary Fig. 6A). Culture of E10.5 3PP explants in the presence of DAPT resulted in the specific and near complete inhibition of mTEC production, evidenced by the absence of UEA1^+^ TEC (Supplementary Fig. 6B, C). In contrast, the numbers of CD205^+^ cTEC/common TEPC were not affected (Supplementary Fig. 6B,C). A few explants contained very rare, isolated UEA1^+^ epithelial cells, and strikingly, these rare K14^+^ or UEA1^+^ TECs were exclusively located in the apparent remnant of 3PP lumen (Supplementary Fig. 6C, arrow), consistent with the localization of CLDN3/4^+^ cells at E10.5 (Hamazaki et al., 2007). Moreover, the number of UEA1^+^ mTECs was unaffected by the presence of RANKL in either control or NOTCH-inhibited conditions (Supplementary Fig. 6B), indicating that the UEA1^+^ epithelial cells present in the cultures represented early, immature mTECs not yet conditioned to respond to thymic crosstalk (Akiyama et al., 2016; Baik et al., 2016).

Collectively, these data unequivocally establish an essential role for that NOTCH-signaling in the normal emergence of the earliest mTEC progenitors, consistent with an obligatory role in mTEC sub-lineage specification. They data further suggest that during normal thymus development, mTEC progenitor emergence commences prior to E12.5.

### Notch activity influences TEC progenitor differentiation

Based on the above data, we wished to test whether NOTCH signaling is permissive or instructive for the specification of mTEC progenitors from the putative common TEPC. We thus developed a TEC-specific NOTCH gain-of-functional model by crossing *Foxn1^Cre^* with *R26-LoxP-stop-LoxP-NICD-IRES-eGFP* (NICD hereafter) mice (Murtaugh et al., 2003), to generate *Foxn1^Cre^;R26-stop-NICD-IRES-eGFP* mice. In this model, high but physiological levels of NICD - and thus constitutively active NOTCH signaling - are heritably induced in all *Foxn1*^+^ cells.

To test whether constitutive NICD expression actively promoted mTEC development, we analyzed TEC differentiation at E14.5, assaying progression of TEC differentiation using PLET1 and MHC Class II (MHCII) as markers of undifferentiated and differentiated cells respectively (Nowell et al., 2011). eGFP expression indicated activation of NICD in >90% of E14.5 TECs (Supplementary Fig. 8; 90.6% ± 1.3%). As expected, a broad down-regulation of PLET1 and up-regulation of MHC Class II (MHCII) was observed in E14.5 control thymi compared to earlier timepoints (Fig. 6A; see Supplementary Fig.1). In contrast, E14.5 NICD thymi exhibited higher proportions of PLET1^+^ and lower proportions of MHCII^+^ TEC than controls. This established that exposure to continuous NOTCH signaling from E12.5 onwards resulted in delayed TEC differentiation (Fig. 6A; see also Supplementary Fig. 1). Analysis of the small population of unrecombined GFP^−^ TEC within the NICD thymus indicated this effect was cell autonomous (Supplementary Fig. 8). The proportion of UEA1^+^ expressing mTECs was unchanged in NICD thymi versus controls, and the cells binding the highest levels of UEA1 were missing (Fig. 6A; NICD, 4.84% ± 0.21%; control 4.43% ± 0.34%) establishing that high NOTCH activity does not drive immediate universal commitment of mTEC at the expense of cTEC.

**Figure 6.**
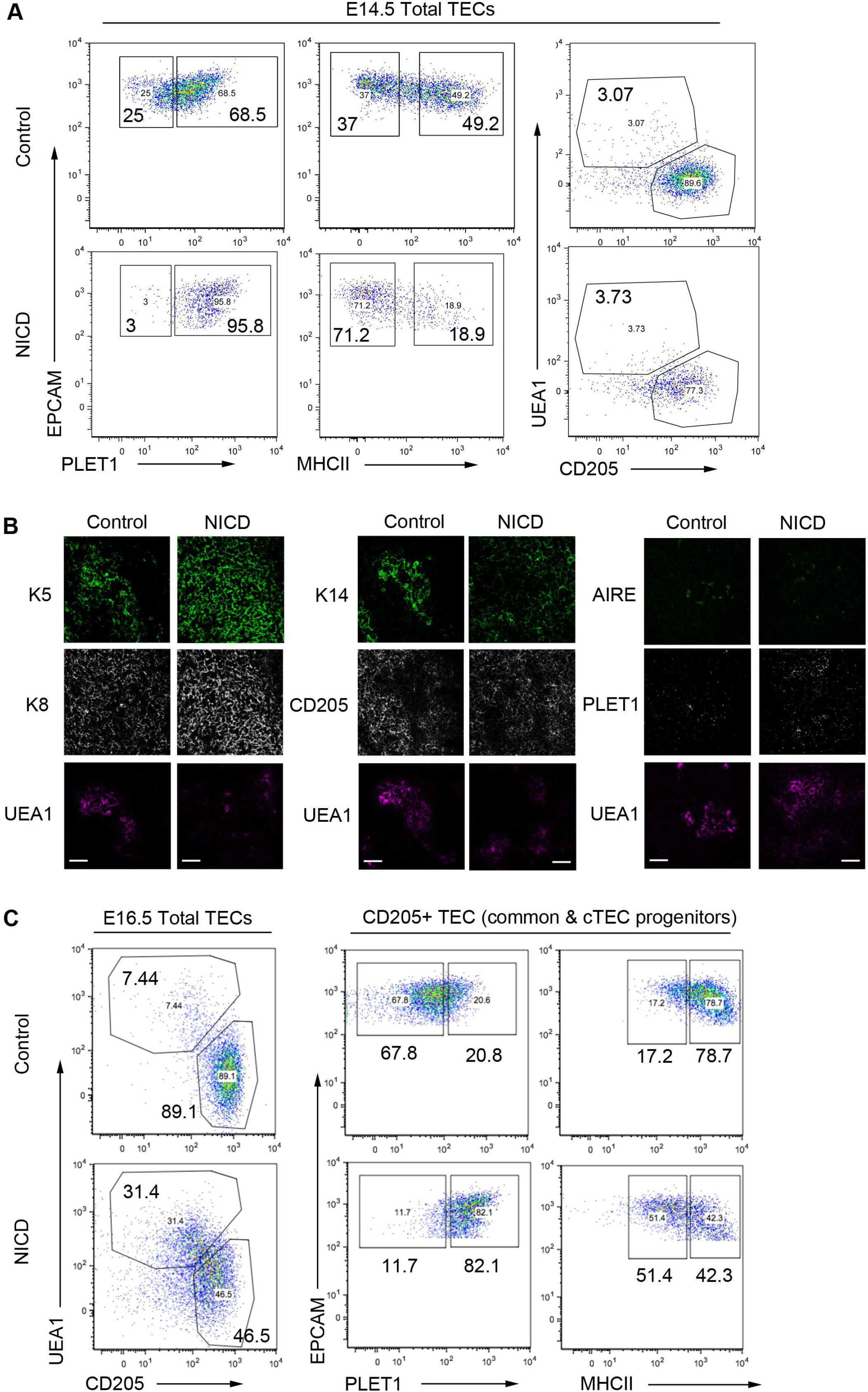
Outcome of enforced NOTCH signaling in TEC. (A) The E14.5 Notch NICD TECs exhibit considerable shift to a PLET1^+^MHCII^−^ immature marker expression. The proportion of UEA1^+^ mTECs is marginally higher in NICD primordia. (B) E16.5 control and NICD thymi stained with the markers shown. (Left) Uniform K5^+^ K8^+^ epithelium in NICD, (middle) expansion of K14 staining into CD205^+^ regions in NICD, compared to clearly demarcated K14^+^ and CD205^+^ zones in controls, (right) both control and NICD thymi express AIRE in UEA1^+^ areas. PLET1 expression is broader in NICD than controls. Scale bar=50μm. (C) (Left) In E16.5 thymi, the percentage of UEA1^+^ mTECs is higher in NICD than in controls. Moreover, more TECs in NICD appear to express intermediate levels of both UEA1 and CD205, or negative for both markers. (Right) The CD205^+^ cTEC/common TEPC population continues to exhibit a PLET1^hi^ MHCII^lo^ immature phenotype. ***Data Collection:*** *Foxn1^Cre^;R26^LSL-NICD-EGFP1^* and C57BL/6 control embryos were collected at E14.5 and E16.5. Samples analyzed were of the same litter. E14.5 NICD, n=4; E14.5 control, n=3; E16.5 NICD, n=3; E16.5 control, n=3. (B) Images are representative of analysis of thymi from two E16.5 NICD and two control embryos.

Since a rapid expansion of mTEC occurs from E14.5, we also analyzed NICD mice at E16.5. These NICD thymi lacked the clearly demarcated medulla present in age-matched controls (indicated by K5, K14 and UEA1). Compartmental boundaries were indistinct, with a pronounced extension of K5 into K8^hi^ CD205^+^ regions suggesting that most TEC had a progenitor cell phenotype (Fig. 6B)(Bennett et al., 2002; Gill et al., 2002; Klug et al., 1998). The NICD sections also contained more extensive PLET1^+^ areas, while containing similar proportions of UEA1^+^ AIRE^+^ mTECs to control thymi. Similarly, flow cytometry analysis showed that the UEA1^+^ and CD205^+^ populations were less clearly defined, with many cells exhibiting an apparently intermediate phenotype (Fig. 6C). Notwithstanding this perturbed distribution, at E16.5 the NICD thymi contained 5-fold more UEA1^+^ mTECs (35.7% ± 7.6%) than control thymi (6.6% ±1.1%). However, the proportion of UEA1^+^ TECs expressing the highest levels of UEA1 was diminished (Fig. 6C). Furthermore, the CD205^+^ cTEC/common progenitors displayed considerably higher PLET1 and lower MHCII levels than controls, consistent with a continued delay/block in cTEC differentiation (Fig. 6C).

Collectively, this establishes that overexpression of Notch promotes but does not dictate mTEC specification from the common TEPC and additionally blocks or substantially delays cTEC lineage progression.

### Impact of NOTCH signaling modulation on gene expression in fetal TECs

To further interrogate the phenotype of NOTCH loss- and gain-of-function models, we analyzed the transcriptome of fetal TECs, aiming to identify mechanisms regulated by NOTCH signaling within specific TEC populations. For both *Rbpj* cKO and control thymi, we collected E12.5 PLET1^+^ TEPCs and E14.5 PLET1^+^ and PLET1^−^ TEC, while for NICD at E14.5 we analyzed only PLET1^+^ TEC, since most NICD TEC were PLET1^+^ at this timepoint (Fig. 6A; for data, see https://www.ncbi.nlm.nih.gov/geo/query/acc.cgi?acc=GSE100314). A trend suggestive of down-regulation of some Notch family and NOTCH target genes was indicated in the E14.5 PLET1 + *Rbpj* cKO versus controls (Supplementary Table 5, Supplementary Fig. 9) and confirmed by RT-qPCR (Supplementary Fig. 10), pointing to a positive feedback loop regulating NOTCH-signaling competence. Conversely, several Notch family genes were significantly upregulated in E14.5 NICD TEC versus controls (Supplementary Table 5, Supplementary Fig. 9).

Independent signaling pathway enrichment analysis using all genes differentially expressed between the E14.5 NICD and wild-type datasets also revealed the NOTCH pathway as one of those most affected by NICD overexpression (Fig. 7A). In addition, we found significant upregulation of the EGFR pathway, known to promote the proliferation of mTEC precursors (Satoh et al., 2016), and of several collagen genes (annotated as “Inflammatory Response Pathway”), suggesting that NOTCH signaling may play a role in endowing proliferative capacity on nascent mTECs and in regulating TEPC differentiation by modifying extracellular matrix (Baghdadi et al., 2018). Neither *Foxn1* nor *Plet1* expression was significantly affected by loss of *Rbpj* (Supplementary Table 5, Supplementary Figs. 9 and 10). The bHLH transcription factor *Ascl1* was markedly down-regulated in *Rbpj* cKO TEC and was also highly enriched in mTEC in wild-type mice, with strong up-regulation occurring co-temporally with medullary expansion at E14.5 (Supplementary Figs. 9, 10 and 11A). This suggested that ASCL1 might act downstream of NOTCH in mTEC lineage regulation. However, no differences in thymic size, organization or cellularity were detected in *Ascl1*^−/−^ thymi (Guillemot et al., 1993) at E17.5 (Supplementary Fig. 11B) apparently ruling out this hypothesis.

**Figure 7.**
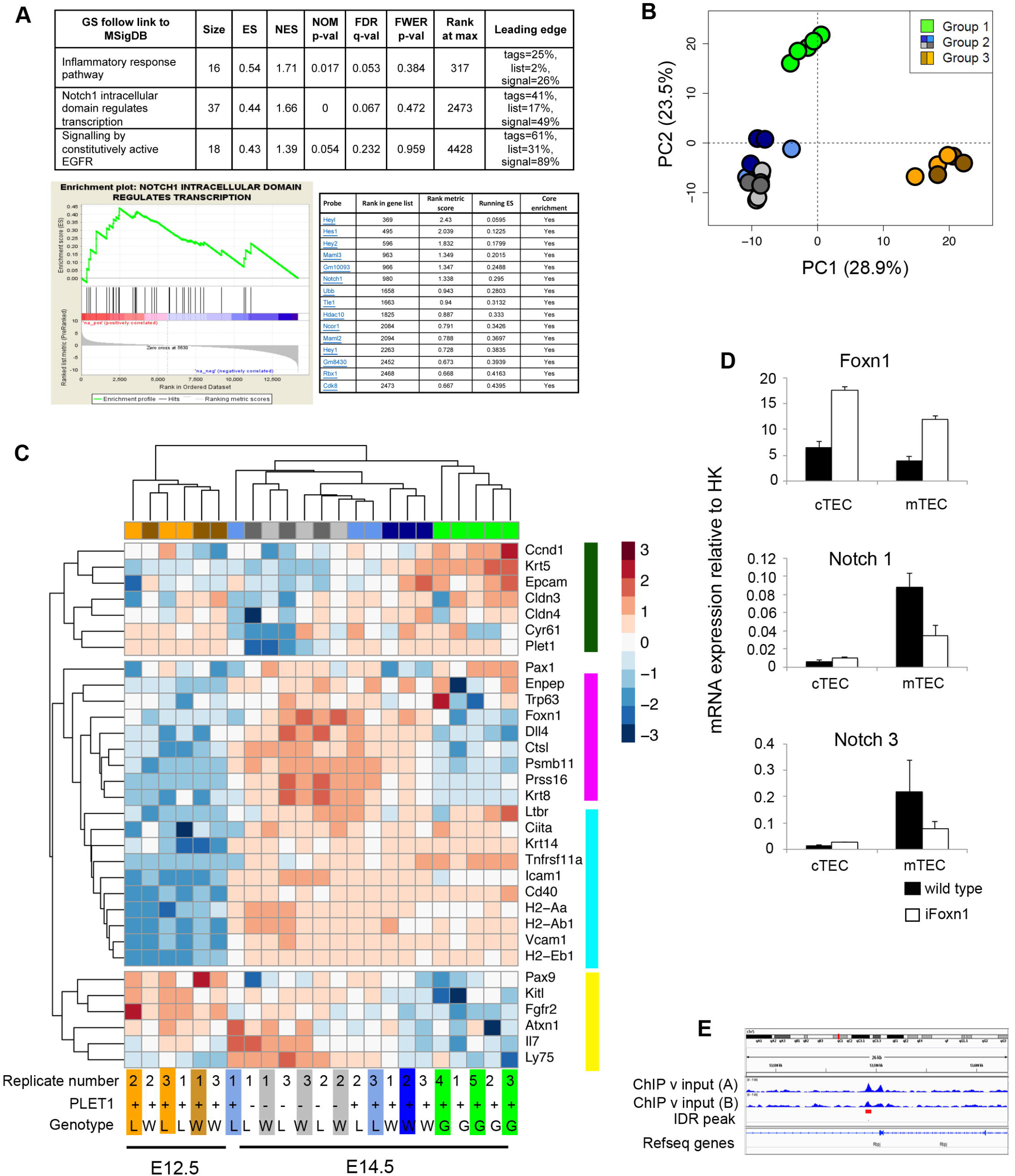
Transcriptome analysis of NOTCH loss and gain of function mutants. **(A)** Pathway analysis of the E14.5 NICD and E14.5 controls identified three signaling pathways as enriched (FDR <= 0.25) in the E14.5 NICD versus the E14.5 controls comparison (top). GSEA enrichment plot for NOTCH signaling pathway (bottom left). Leading edge subset genes contributing to the enrichment for NOTCH signaling pathway (bottom right). **(B)** PCA of *Rbpj* cKO, wild type and NICD TEC at the ages shown (500 most variable genes). Group 1, E14.5 NICD samples; Group 2, E14.5 PLET1^+^ and PLET1^−^ *Rbpj* cKO and controls; and Group 3, E12.5 *Rbpj* cKO and controls. **(C)** Heatmap of lineage specific genes among all groups of samples shown in PCA above. Colors on top and bottom of the heatmap indicate clustering of samples per group, while side colors indicate groups of genes regulated similarly across conditions. Groups: E12.5 wild type, brown; W12.5 *Rbpj* cKO, orange; E14.5 wild type PLET1^+^, dark blue; E14.5 wild type PLET1^−^, light grey; E14.5 *Rbpj* cKO PLET1^+^, light blue; E14.5 *Rbpj* cKO PLET1^−^, dark grey; W, wild type; L, loss of function (*Rbpj* cKO); G, gain of function (NICD). **(D)** RT-qPCR analysis of sorted cTEC and mTEC from E17.5 wild-type and iFoxn1 thymi for the genes shown. Error bars show mean±SD. **(E)** Genomic locus of *Rbpj* showing Foxn1 peaks identified in (Zuklys et al., 2016). ***Data Collection:*** (A-C) To obtain the E12.5 and E14.5 cKO and wild type samples, thymi were microdissected from E12.5 and E14.5 embryos generated from a *Foxn1^Cre^;Rbpj^FL/+^ x Rbpj^FL/FL^* cross and TECs obtained by flow cytometric cell sorting. Following genotyping, cells from three cKO and three control samples were processed for sequencing. The E12.5 and E14.5 samples were each obtained from two separate litters, on two separate days for each timepoint. To obtain the E14.5 NICD samples, thymi were microdissected from five E14.5 Foxn1Cre; R26^LSL-NICD-EGFP^ embryos of the same litter, TECs were obtained by flow cytometric cell sorting, and the samples processed for sequencing. (D) n=3, where each n represents TECs sorted from pooled embryos from a single litter of E17.5 iFoxn1 or wild-type embryos.

Principal Component Analysis (PCA) clustered the E12.5 and E14.5 PLET1^+^ *Rbpj* cKO and wild type, and E14.5 PLET1^+^ NICD, datasets into three groups, E14.5 NICD samples (Group 1); E14.5 PLET1^+^ and PLET1^−^ *Rbpj* cKO and controls (Group 2; see also Supplementary Fig. 12); and E12.5 *Rbpj* cKO and controls (Group 3)(Fig. 7B). The broad PCA analysis (Fig. 7B) separated the samples by developmental stage (PC1) and PLET1 level (PC2; note that PC2 is not solely PLET1), with Group 1 positioned between Group 2 and Group 3 in PC1. Overall, the PCA is consistent with E14.5 NICD TEC exhibiting at least a partial developmental delay (in keeping with conclusions from Fig. 6), or sustained NICD expression in early TEC inducing a distinct cell state that is not found/ is very rare in the early wild type fetal thymus.

Consistent with these possibilities, clustering analysis revealed differential effects of NOTCH signaling perturbation on markers associated with differentiation into the cTEC and mTEC sublineages, general TEC maturation, or the earliest TEPC state. In particular, genes associated with cTEC lineage identity (*Ctsl, Dll4, Psmb11, Prss16, Krt8, Ly75*) were up-regulated normally from E12.5 to E14.5 in the *Rbpj* cKO samples but were expressed at levels similar to E12.5 wild-type in the E14.5 NICD samples (Fig. 7C), consistent with maintained NOTCH signaling imposing a block on cTEC generation from the common TEPC/early cTEC progenitor. *Foxn1* also exhibited this expression pattern (Fig. 7C), and indeed many genes in this panel are direct FOXN1 targets (Calderon and Boehm, 2012; Nowell et al., 2011; Zuklys et al., 2016). Notably, overexpression of FOXN1 led to down-regulation of a number of NOTCH family and NOTCH target genes in fetal TEC (Fig. 7D and data not shown), suggesting that induction of FOXN1 may down-regulate NOTCH signaling in TEC during normal development *in vivo*. Consistent with this, our reanalysis of published FOXN1 ChIP-seq data (Zuklys et al., 2016) indicated *Rbpj* as a direct FOXN1 target (Fig. 7E). Moreover, Zuklys and colleagues (Zuklys et al., 2016) identified several known NOTCH targets and modulators as FOXN1 targets (*Hey1, Hes6, Deltex4* and *Fbxw7)*. The relative down-regulation of *Foxn1* resulting from sustained NICD expression in early fetal TEC (Fig. 7C, Supplementary Fig. 9) thus suggests the possibility of reciprocal inhibition.

Other genes associated with both cTEC and mTEC differentiation, were unaffected or only marginally affected by the NOTCH signaling gain- or loss-of-function mutations. In contrast, markers associated with the mTEC sub-lineage (*Krt5, Epcam*) were strongly up-regulated in the E14.5 NICD samples compared to controls, and these genes also clustered with other genes normally strongly down-regulated from E12.5 to E14.5 (*Cldn3, Cldn4, Cyr61, Plet1, Ccnd1*). Tnfrs11a, the gene encoding RANK, was also significantly up-regulated in the E14.5 NICD versus wild type samples (Fig. 7C). Finally, a category including *Pax9, KitL* and *Fgfr2*, which are normally highly expressed at E12.5, was markedly down-regulated in the E14.5 NICD compared to other E14.5 samples (Fig. 7C).

Overall, we conclude that upregulation of NOTCH signaling in TEC during early thymus development at least partially blocks cTEC differentiation and promotes but does not dictate mTEC development, suggesting that NOTCH regulates not only mTEC specification but also maintenance of the fetal thymus common TEPC (Fig. 8).

**Figure 8.**
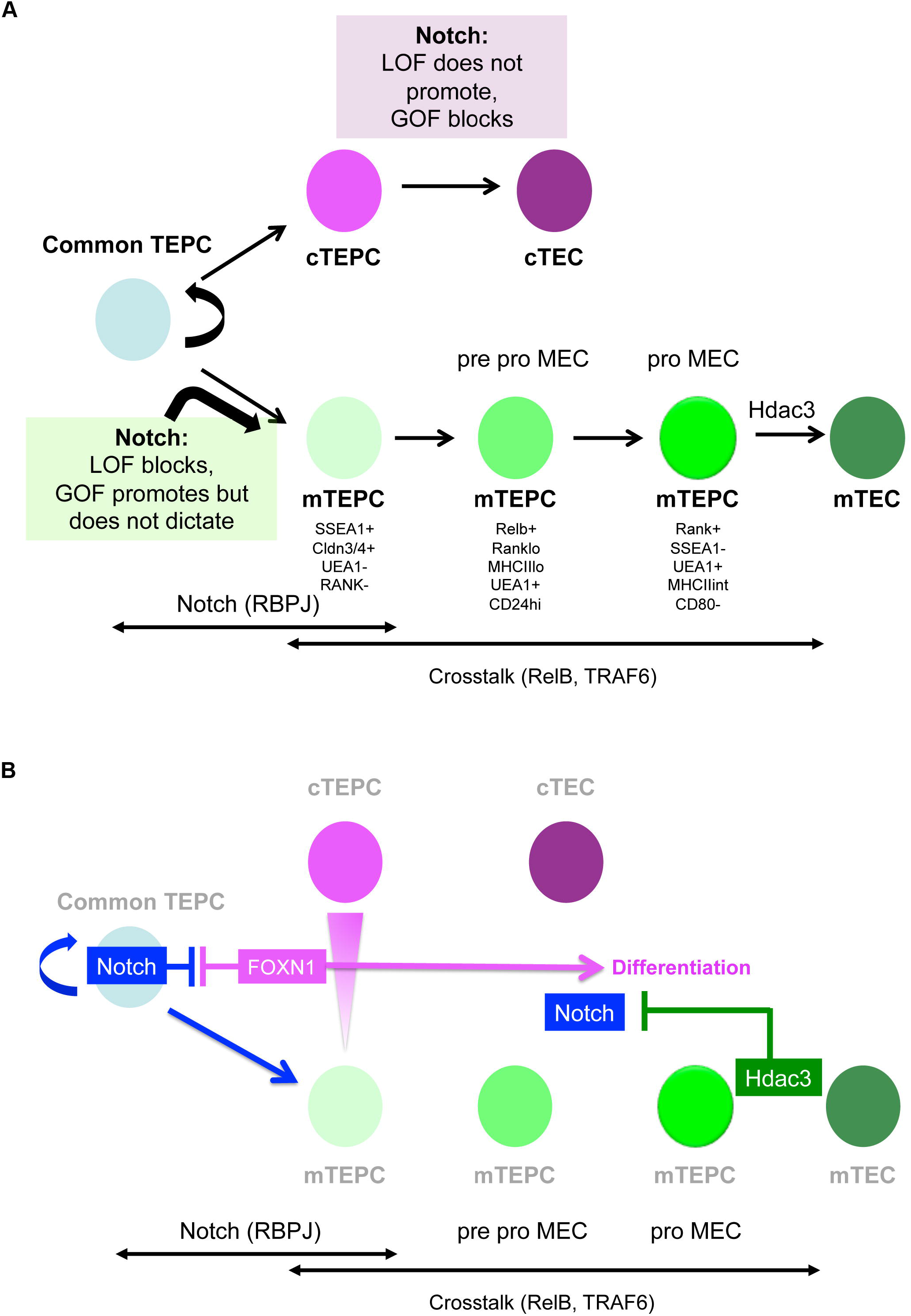
Model for NOTCH signaling regulation of early TEC development.

## Discussion

We have used conditional loss- and gain-of-function approaches together with pharmacological inhibition to investigate the role of NOTCH signaling in TEC. Our data show, based on TEC-specific RBP-Jκ deletion, γ-secretase inhibition in FTOC and enforced dnMAML expression in the developing endoderm from E9.5, that NOTCH activity is essential for mTEC development. Specifically, they establish that NOTCH signaling is required for the emergence of the mTEC sub-lineage from the putative bipotent TEC progenitor, strongly suggesting that NOTCH regulates mTEC specification, and further show that during mouse fetal thymus development, this requirement is restricted to a developmental window prior to E16.5. Additionally, they demonstrate that NOTCH signaling, whilst essential, is permissive rather than instructive for mTEC development and indicate a further role for NOTCH in regulating exit from the early bipotent TEPC state into mTEC and cTEC differentiation. These findings, summarised schematically in Fig. 8, raise several issues which are discussed below.

### Timing of the NOTCH signaling-requirement

NOTCH signaling has been shown to regulate distinct events in the different developmental stages of a tissue (Hartman et al., 2010; Radtke et al., 2004; Shih et al., 2012). A recent study reported that NOTCH activity is enriched in cTECs and that repression of NOTCH by the histone deacetylase HDAC3 is important for expansion/maintenance of developing mTEC (Goldfarb et al., 2016). This study analyzed the same NOTCH overexpression line as used herein, but at the later time-points of 10 days and 6 weeks postnatal (Goldfarb et al., 2016). The conclusions of this and our own studies are entirely compatible, with the data presented herein establishing a requirement for NOTCH signaling at the earliest stages of TEC lineage divergence, and the data of Goldfarb indicating that down-regulation of NOTCH signaling is required for later stages of mTEC differentiation (Goldfarb et al., 2016). Indeed, the phenotypes observed in each may reflect the outcome of same perturbation in TEC differentiation, at separate stages. However, it is also possible that NOTCH has secondary roles in TECs subsequent to its initial role in mTEC specification, which ceases by E16.5 (Fig. 2).

### Thymic crosstalk

The NFκB pathway plays a vital role in mTEC development and consequently in the establishment of central tolerance (Akiyama et al., 2005; Burkly et al., 1995; Kajiura et al., 2004). Recent studies using transcriptomics and functional assays have led to more clarity on how the several NFκB ligands, through which thymic crosstalk occurs, function during mTEC maturation (Akiyama et al., 2016; Bichele et al., 2016; Desanti et al., 2012; Mouri et al., 2011). In particular, Akiyama and colleagues identified two separable UEA1^+^ mTEC progenitor stages, pro-pMECs and pMECs based on the expression of RANK, MHCII and CD24 (Akiyama et al., 2016). The transition from the more primitive pro-pMECs to pMECs depends on RELB, whereas further maturation from pMECs is TRAF6-dependent. Crucially, both pro-pMECs and pMECs respond to induction by RANKL in T cell-depleted FTOC (Akiyama et al., 2016). We initially interpreted our data on potential interplay between NOTCH and NFκB to suggest synergy between these two pathways in mTEC development, since NOTCH signaling-inhibition attenuated RANKL stimulation in E15.5 wild-type T cell-depleted FTOC (Fig. 4A). However, comparison with E15.5 *Rbpj* cKO FTOC indicated that NFκB activation of already-specified mTEC progenitors is unaffected by lack of NOTCH signaling-responsiveness: although the block in mTEC development was more severe in the *Rbpj* cKO FTOC the few mTEC that were present could be stimulated by RANK, indicating the presence of pMECs and/or pro-pMECs. The attenuation of RANKL stimulation upon DAPT treatment of E15.5 wild-type FTOC thus suggests that mTEC specification is still on-going at E15.5. However, we also observed that mTEC clusters in *Rbpj* cKO thymi tended to be smaller than those in controls, and therefore the possibility that in addition to regulating mTEC specification NOTCH also regulates the initial expansion of mTEC progenitors cannot be ruled out. Indeed, our data reveal EGFR signaling as a major target of NOTCH during early TEC development.

In contrast, our data show that while E10.5 3PP explants can generate UEA1^+^ mTECs and CD205^+^ cTEC/progenitors in culture, these UEA1^+^ mTEC do not respond to RANKL. It is thus likely that the UEA1^+^ cells in these explants represent an even more primitive mTEC progenitor state than the pro-pMECs. Of note is that some DAPT-treated E10.5 3PP explants produced no UEA1^+^ mTECs, and thus that mTEC specification can be completely suppressed in the absence of NOTCH signaling. Taken together, these results suggest that although NOTCH and NFκB are both required for mTEC development, the two pathways act sequentially but independently.

### Notch regulation of mTEC progenitor emergence

The loss of mTECs in NOTCH loss-of-function models could be explained by three hypotheses: (i) NOTCH might regulate the decision of bipotent TEPCs to become mTECs. In this model, in the absence of NOTCH signaling, bipotent progenitors fail to commit to mTEC fate and over time become cTECs instead. (ii) Alternatively, high levels of NOTCH signaling might dictate that TEPCs remain bipotent, with cells that experience lower NOTCH committing to the cTEC lineage. Unlike the ‘specification hypothesis’, in this scenario mTECs would fail to emerge in the absence of NOTCH signaling because the bipotent TEPCs undergo premature differentiation into cTECs, exhausting the pool that retains the potential for mTEC generation. (iii) Finally, NOTCH might be required for the proliferation of specialized mTEC progenitors; in this case we would expect the perturbation to affect only mTECs and not cTECs or bipotent progenitors.

We conclude from the gain-of-function data that enhanced NOTCH activity neither switches all TECs to become mTECs, nor only affects mTECs. Instead, NOTCH activity is necessary but not sufficient for mTEC fate in the developmental timeframe investigated. Despite the caveats with established markers, the considerable shift towards a PLET1^+^MHCII^−^ (Fig. 6A, C) K5^+^ K8^+^ (Fig. 6B) phenotype suggests a more immature, TEPC-like state as the primary phenotype resulting from high NOTCH activity. Indeed, the transcriptome of E14.5 NICD TECs occupies a state that is separate from both E12.5 TEPCs and age-matched controls, whilst sharing certain features with both clusters. As development progresses from E14.5 to E16.5, many TECs do upregulate the mTEC markers UEA1 and K14, indicating that high NOTCH activity is compatible with acquisition of mTEC fate. Importantly, the NICD^+^ UEA1^+^ mTECs at E16.5 display comparable maturation status to controls, whereas CD205^+^ cTEC/common TEPCs continue to exhibit a primitive phenotype (Fig. 6). These data suggest that once mTECs are specified, further development is independent of NOTCH signaling.

The gain-of-function results also support our hypothesis that NOTCH operates at the TEC progenitor level, whilst opposing the model that NOTCH activity only influences mTECs. It does not however rule out the specification model. Although retention of an early progenitor state seems to be the primary outcome of enforced NOTCH signaling, the proportion of mTEC in the E16.5 gain-of-function thymi is higher than controls. Several factors may be in play in this second phase. The duration of signaling has been shown to result in the temporal adaptation of sensitivity in several pathways (reviewed in (Kutejova et al., 2009)). Moreover, instead of a simple ON/OFF response, the NOTCH response may be graded, as in the case of inner ear (Petrovic et al., 2014) and pancreas development (Shih et al., 2012). mTEC specification may require higher levels of NOTCH, which could for instance be achieved by positive feedback above the levels of those imposed by the enforced NICD expression in the NICD hemizygous mice used in these experiments. Variables independent from NOTCH may also play a part. A potential candidate is FOXN1, which drives TEPCs out of the primitive undifferentiated state and into differentiation (Nowell et al., 2011), and indeed our data indicate interplay between FOXN1 expression levels and NOTCH activity (as depicted in Fig. 8). In addition to the direct interaction suggested from our analysis, FOXN1-mediated repression of NOTCH activity could be reinforced via its direct targets DLL4 and FBXW7; the former may mediate cis-inhibition of NOTCH receptors, while the latter has been shown to enhance the degradation of NICD (Carrieri and Dale, 2016; del Alamo et al., 2011). We note that the thymic phenotype of the NOTCH gain-of-function mutant reported here resembles those of the *Foxn1^R/−^* (Nowell et al., 2011) and the *Foxn1^Cre^;iTbx1* (Reeh et al., 2014) mutant mice, in which exit from the earliest TEPC compartment is also severely perturbed due to the inability to express normal levels of FOXN1.

One of the long-term goals of the field is to create fully functional thymus organoids from TECs derived from pluripotent stem cells or by direct conversion from unrelated cell types (reviewed in (Bredenkamp et al., 2015)). Understanding the duration of TEPC bipotency, lineage plasticity and NOTCH activity would improve protocols and inform strategies in this regard. Our data predict that, by manipulating the levels of NOTCH signaling TEPCs experience, it may be possible to produce more homogenous populations of TEC subsets, including TEPC. However, the complexities indicated from studies on NOTCH in other organs, together with the potential for differential effects on TEC at different stages of lineage progression, suggest that further advances in this direction will require caution and precision.

## Material and methods

### Mice

CBAxC57BL/6 F1 mice were used for isolation of fetal TEC. For timed matings, C57BL/6 females were housed with CBA males, and noon of the day of the vaginal plug was taken as E0.5. Representative data shown were obtained from littermates or, when not possible, embryos sharing the same plug date. *Foxn1^Cre^* (Gordon et al., 2007), *Rbpj* conditional knockout (Han et al., 2002), *Rosa26-stop-NICD* (Murtaugh et al., 2003), *CBF1*-Venus (Nowotschin et al., 2013), *Ascl1*^−/−^ (Guillemot et al., 1993), *Rosa26^CreERt2/CAG-Foxn1-IRES-GFP^* (iFoxn1) (Bredenkamp et al., 2014), and *Foxa2^T2iCre^;Rosa26^loxp-STOP-loxp-dnMAML-IRES-eGFP^* and *Foxa2^T2iCre^;Gt(ROSA)26So^rtm1(EYFP)Cos^* (Horn et al., 2012; Maillard et al., 2004) mice were as described. All animals were housed and bred at the CRM animal facilities except for the *Ascl1*^−/−^ strain, which was housed and bred at NIMR, Mill Hill, London; the *Rosa26NICD* strain (Murtaugh et al., 2003), which was housed and bred at EPFL, Lausanne; and the *Foxa2^T2iCre^, Rosa26^loxp-STOP-loxp-dnMAML-IRES-eGFP^* (Horn et al., 2012; Maillard et al., 2004) and *Gt(ROSA)26Sor^tm1(EYFP)Cos^* (R26LSL-YFP) (Srinivas et al., 2001) strains which were housed and bred at DanStem, University of Copenhagen. *Foxn1^Cre^* (Gordon et al., 2007) were also housed and bred at EPFL, Lausanne. All experimental procedures were conducted in compliance with the Home Office Animals (Scientific Procedures) Act 1986.

### Thymus dissociation

Postnatal thymi were dissociated in 1.25mg/ml collagenase D (Roche), and subsequently in 1.25mg/ml collagenase/dispase (Roche) diluted in RPMI medium (Life Technologies). 0.05mg/ml DNaseI (Lorne) was added to the buffer to minimize cell adhesion. Fetal thymi were dissociated for 20 minutes using a PBS-based buffer consisting of 1.25mg/ml collagenase D, 1.4mg/ml hyaluronidase (Sigma) and 0.05mg/ml DNaseI. After digestion cells were spun down and digested in 1x trypsin for two minutes. Cell suspension was then filtered through 70μm cell strainer (Corning) to remove clumps.

### Flow Cytometry

Adult thymi and grafted RFTOC were processed for flow cytometric sorting and analysis as previously described (Bredenkamp et al., 2014; Nowell et al., 2011). See Supplemental Experimental Procedures for detailed protocols. For analysis and sorting, adult thymic tissue was depleted of T cells using anti-CD45 MACS beads (Miltenyi Biotec); fetal tissue was not T cell depleted. Cell counts were carried out using a BioRad cell counter and slides, where required. Sorting and analysis was performed using a BD FACS Aria II and a BD LSR Fortessa respectively at the CRM, University of Edinburgh. For *Rosa26NICD* TEC, sorting was performed on a BD FACS Aria II at the University of Lausanne, Epalinges. Sorting protocols were identical for all cell isolation experiments. All flow cytometry data were analyzed using FlowJo Version 9.7.6 (Tree Star, Inc).

### Immunohistochemistry

Immunohistochemistry was performed as described (Gordon et al., 2004). See Supplemental Experimental Procedures for details. Appropriate isotype and negative controls were included in all experiments. For detection of immunofluorescence, slides were examined with Leica SP2, SPE and SP8 (Leica Microsystem, GmbH) confocal microscopes. Images presented are of single optical sections. Fiji software (Schindelin et al., 2012) was used to quantify the surface area of positive staining and the thymic section. Volume percentage of K14^+^ or UEA1^+^ regions in an embryo was defined as total area of positive staining divided by total area of thymic section.

### Antibodies

The antibodies used for immunohistochemistry and flow cytometry were as listed in Table S2. See also Supplemental Experimental Procedures.

### Medium

TEPC medium was N2B27 (DMEM) medium, 20ng/ml BMP4, 20ng/ml FGF8, Penicillin/streptomycin, 1μg/ml heparin.

### Fetal thymus organ culture (FTOC)

For reaggregates or thymi older than E15, FTOCs were cultured on a Millipore membrane raft floating on DMEM supplemented with 10% FCS and L-glutamine. Third pharyngeal pouches or thymic primordia younger than E15.5 were submerged and allowed to settle on thin matrigel (Corning), then cultured in N2B27 supplemented with BMP4 (Peprotech) and FGF8 (Peprotech). Where DAPT (Tocris) or deoxyguanosine (dGUO; Sigma) were used, the equivalent amount of DMSO was added to the control medium. RANKL (Peprotech) was used at 500ng/ml.

### Quantitative real-time PCR

RT-qPCR was performed as previously described (Bredenkamp et al., 2014), on 50-200 cells per sample. See Supplemental Experimental Procedures for details. Data are shown after normalization to the geometric mean of three control genes (*Hprt, Ywhaz, Hmbs*). Data analysis was carried out using LightCycler 1.5 software and the ΔCt method (Livak and Schmittgen, 2001). Primers used for RT-qPCR are as shown in Table S3.

### RNA-seq

100 cells were sorted directly into Smartseq2 lysis buffer (Picelli et al., 2013) at the CRM, University of Edinburgh (*Rbpj* cKO and littermate control samples) or at the University of Lausanne, Epalinges, Ch (*Rosa26NICD* samples). Sorted samples were immediately frozen on dry ice and were then shipped to the WIMM, University of Oxford for library preparation. The libraries were then prepared and sequenced at the Wellcome Trust Centre for Human Genetics, University of Oxford. Quality control (QC) of the raw reads by FastQC (Andrews, 2010) indicated small amount of adaptor contamination and few low quality reads, therefore the raw data were trimmed with Trimmomatic (Bolger et al., 2014) using default parameters for PE reads and the cropping option specific for the Nextera PE adapters. Only paired reads that passed QC were aligned with STAR against the mouse genome assembly (GRCm28 – Ensembl 87) and the aligned reads were assigned to genes with featureCounts (Liao et al., 2014). The resulting count tables were imported to R for further normalisation and analysis. Batch effect correction was applied for the within group lane effects, however, some batch effects could not be corrected. This applied to the potential for a laboratory effect between the E14.5 NICD and all other samples, since the E14.5 NICD sample was collected at EPFL Lausanne. However, the same thymus dissociation and cell sorting protocols, and the same make and model of cell sorter, were used, and the subsequent sample processing was performed at the University of Oxford using the same protocol as for all of the other samples. To control for this, the expression levels of housekeeping genes were determined for all samples and were not biased in any particular groups (Supplementary Fig.12B).

Differential expression analysis was performed using the LIMMA package and voom (Ritchie et al., 2015) from Bioconductor (Gentleman et al., 2004) and a threshold of FDR ≤ 0.05 was set to define genes that change with significance between the different datasets. The table of all differentially expressed genes and their fold changes was used as a pre-ranked list in GSEA (Subramanian et al., 2005) against the ConsensusPathDB (Kamburov et al., 2011) to predict signaling pathways that are enriched between the wild type and NICD samples. Pathways were defined as enriched if they had an FDR value ≤ 0.25 (default significance criteria for GSEA).

### ChIP-seq

Publicly available data under GEO accession number: GSE75219 (Zuklys et al., 2016) were reanalyzed as follows. QC of the raw reads by FastQC (Andrews, 2010) indicated a few low quality reads, and these were therefore removed trimming the raw data with Trimmomatic (Bolger et al., 2014) using default parameters for PE reads. Read mapping was performed with Bowtie2 (Langmead and Salzberg, 2012) with default parameters; MACS2 (Zhang et al., 2008) was used with a lenient p-value threshold of 1×10^−3^ to call peaks. The IDR pipeline (Li et al., 2011) was followed to call confident peaks among replicates (IDR<=0.05).

### Data availability

RNAseq data are deposited in GEO, and are available through the following link: https://www.ncbi.nlm.nih.gov/geo/query/acc.cgi?acc=GSE100314.

### Statistics

Statistical analysis was performed using the GraphPad Prism 7.02 software. Student’s t-test (two-tailed, unpaired) was performed for pair-wise comparisons. Multiple comparison procedures were performed with one-way ANOVA test (two tailed), as appropriate for normally distributed data (normal distribution was tested using Chi2 goodness of fit). The alpha level is taken as 0.05. Errors shown are standard deviations (s.d.) throughout. Sample sizes of at least n=3 were used for all analyses except where indicated. For all analyses, n represents the number of independent biological experiments. No statistical method was used to predetermine sample size, the experiments were not randomized, and the investigators were not blinded to allocation during experiments and outcome assessment. There were no limitations to repeatability of the experiments. No samples were excluded from the analysis.

### Author Contributions

DL and AIK conceived and designed experiments, performed experiments, analyzed the data and contributed to writing the manuscript; KEO’N, AMF and MP conceived and designed experiments, performed experiments and analyzed data; UK performed experiments and analyzed data; SRT, SU, PS, UK, FR, PS and FG contributed to experimental design and analysis of the data; CCB conceived the original idea, designed experiments, contributed to analysis of the data and wrote the manuscript. The authors declare no competing financial interests.

## Acknowledgements

We thank C. Cryer and F. Rossi (CRM, University of Edinburgh), and R. Bedel, A. Ribeiro and A. Wilson (FCF UNIL, Lausanne), for cell sorting, the Biomed Unit staff for animal care. The research leading to these results received funding from the School of Biological Sciences, University of Edinburgh (DL), the Medical Research Council (CCB), the Biotechnology and Biological Sciences Research Council (BB/H021183/1BBSRC; KO’N, CCB), the European Union Seventh Framework Programme (FP7/2007-2013) collaborative projects EuroSyStem (CCB, FR and UK), OptiStem (CCB, FR and UK) and ThymiStem (CCB, SRT, DL, AIK, KO’N) under grant agreement numbers 200720, 223098 and 602587 respectively.

